# Hair Cells Loss Estimation from Audiograms

**DOI:** 10.1101/2024.07.07.602394

**Authors:** Miriam Furst, Yonatan Koral, Asaf Zorea

## Abstract

Age-related hearing loss is characterized by a progressive loss of threshold sensitivity, especially at high frequencies. There is increasing evidence that the loss of cilia in the inner and outer hair cells is the dominant cause of hearing loss. We present a framework for calculating the human auditory threshold based on a non-linear time-domain cochlear model that incorporates hair cell damage along the cochlear partition. We successfully predicted the audiogram measured prior to death by substituting the postmortem percentage of surviving hair cells, using data from Wu et al. (Wu *et al*., 2020). We also present an algorithm for estimating the percentage of hair cells from a measured audiogram. Comparison with the data from Wu et al. revealed that the algorithm accurately predicted the surviving inner hair cells along the entire cochlear partition and the outer hair cells at the basal part of the cochlea.

## I. INTRODUCTION

Age-related hearing loss (presbycusis) is a common phenomenon occurring in modern society. It is characterized by a progressive loss of threshold sensitivity, especially at high frequencies. Accurate classification of pathologies that cause the hearing loss (HL) could improve our understanding of hearing mechanisms, HL diagnosis, and treatment strategies. Identifying the multiple contributors to the audiometric loss of a hearing-impaired listener remains an ongoing challenge (Dubno *et al*., 2013; Jepsen and Dau, 2011; Johannesen *et al*., 2014; Moore, 2004; Moore and Glasberg, 1997; Plack *et al*., 2004; Schmiedt, 1996; Vaden *et al*., 2022).

The most common cochlear pathologies include hair cell damage due to loss of cilia, degeneration of the stria vascularis affecting the endocochlear potential in the scala media, and auditory nerve degradation. Several approaches exist for estimating cochlear pathology from audiogram patterns. Dubno et al. (Dubno *et al*., 2013) defined two main phenotypes of impaired audiograms: the metabolic phenotype resulting from the deterioration of the stria vascularis in the cochlear lateral wall, and the sensory phenotype, resulting from damage to sensory cells in the inner ear, particularly the outer hair cells (OHC). Each individual impaired audiogram is estimated as a combination these phenotypes, which eventually indicate the possible pathologies causing HL (Dubno *et al*., 2013; Schmiedt, 2010; Vaden *et al*., 2022). Another approach assumes that the hearing loss of any test frequency results from cochlear mechanical gain loss, or specifically OHC dysfunction and inefficient inner hair cell (IHC) transduction. The contribution of IHC and OHC to hearing loss is determined from the cochlear I/O curves, which were obtained by measuring the temporal masking curve (Johannesen *et al*., 2014; Moore and Glasberg, 1997; Plack *et al*., 2004).

Up to this point, evaluating the different methods has been challenging due to lack of direct measurements of threshold shifts and cochlear pathology in humans. Recent histopathological analysis of normal-aging human cochleae, whose audiograms were obtained close to the time of death (Wu and Liberman, 2022; Wu *et al*., 2020) provides an opportunity to test the ability to predict cochlear pathology from individual audiograms. Their statistical analysis indicated that damage to the hair cells along the cochlear partition, particularly due to missing cilia, predicts audiometric shifts better than neural loss or trial atrophy.

This study conducts a comparative analysis using data from Wu et al. (Wu *et al*., 2020) and an end-to-end computational model of the peripheral auditory pathway (Faran and Furst, 2023; Furst, 2015). The model comprises a time-domain non-linear cochlear model with integrated OHC functionality, a model of IHC transduction process, and an optimal estimation of the auditory threshold. An impaired auditory threshold can be identified as a result of damage to IHCs and OHCs along the cochlear partition.

The current study has two primary objectives based on the computational model: (1) to assess whether post-mortem hair cell loss along the cochlear partition can predict individual audiograms and (2) to develop an algorithm capable of estimating hair cell loss from audiometric measurements.

## II. THE COMPUTATIONAL MODEL

The computational model is schematically described in Fig. 1. A detailed description of the model is given in Appendix. A. The input to the model is an acoustic signal, and its output is the estimated auditory threshold. For sinusoidal inputs, the output is the estimated audiogram. The process begins with the middle ear and the cochlea. The involvement of cochlear sensory cells is specifically represented by two parameters: *γ*_ohc_(*x*) and *γ*_*ihc*_(*x*) for the OHCs and IHCs, respectively. The OHCs modify the basilar membrane (BM) displacement through electromotility feedback, while the IHCs transduce the BM motion into neural signals for the auditory nerve.

**FIG. 1.**
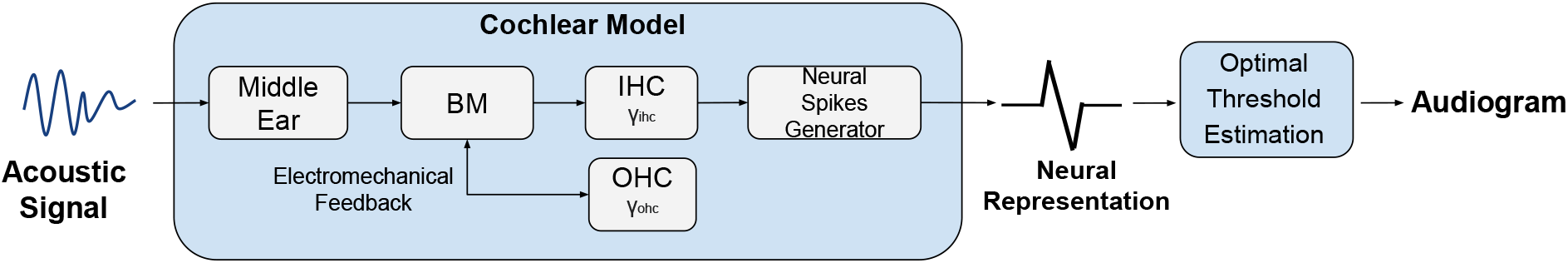
Model Outline

### A. Audiogram Estimation as a Function of Hair Cell Loss

This section presents a model for predicting the auditory threshold given the percentage of survived hair cells along the cochlear partition.

The contribution of the OHCs to the BM motion is expressed by the following equation (Barzelay and Furst, 2011; Cohen and Furst, 2004; Furst, 2015):

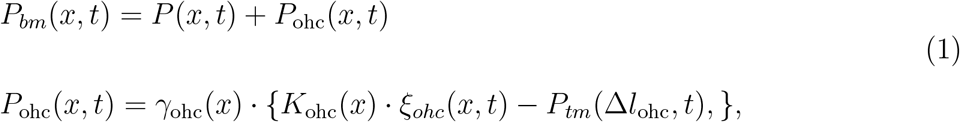

where *P*_*bm*_ and *P*_*tm*_ are the mechanical pressures of the BM and tectorial membrane (TM), respectively. *P* represents the differential fluid pressure between scala tympani and scala vestibuli. *ξ*_*ohc*_ = *ξ*_*tm*_ −*ξ*_*bm*_ indicates the displacement of the OHC, derived from the difference between the *TM* (*ξ*_*tm*_) and *BM* (*ξ*_*bm*_) displacements. Δ*l*_*ohc*_ denotes the OHC length change obtained by the electromotility process (Brownell *et al*., 1985). *K*_ohc_ denotes the OHC stifness. *γ*_ohc_ represents the effectiveness of the OHCs, reflecting the number of cells and cilia in each cell.

The contribution of the IHCs is expressed via their electrical voltage, *ψ*_*ihc*_, as follows (Furst, 2015; Zilany *et al*., 2014, 2009):

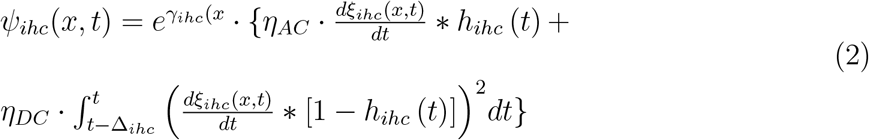

where *h*_*ihc*_(*t*) is the impulse response of the IHCs, *ξ*_*ihc*_ ≈ *ξ*_*bm*_. *η*_*AC*_, *η*_*DC*_, and Δ_*ihc*_ are constants. *γ*_*ihc*_ represents the efficiency index of the IHCs, which correlates to the number of cilia in each cell. The instantaneous rate of auditory nerve spiking is then obtained by the following equation:

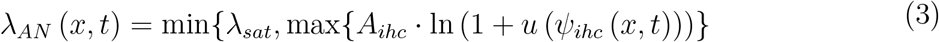

where *λ*_*sat*_ is the saturation rate of the auditory nerve, *u*(*t*) is a step function, and *A*_*ihc*_ is constant.

The auditory threshold can be derived by either the average auditory nerve response or the synchronous response (Heinz *et al*., 2001; Siebert, 1970). Based on our previous study (Furst, 2015), we utilized the average rate response, which has shown that it can predict the auditory threshold in the presence of noise, whereas the synchronous response tends to yield unrealistically low thresholds. Since the neural response is modeled as a non-homogeneous-Poisson-process (NHHP), the auditory threshold can be obtained using the Cramer-Rao Lower Bound (CRLB), which yields the amplitude threshold for the *m*^*th*^ fiber

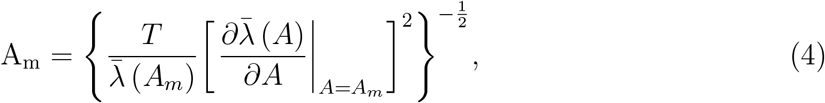

where for every frequency (*f*) the input stimulus is *s*(*t*) = *Asin*(2*πft*), *T* is the stimulus duration, *A* is the stimulus amplitude, and 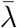 is the average rate of response. Based on the entire auditory nerve comprising *M* = 30, 000 single fibers, the auditory threshold is calculated as:

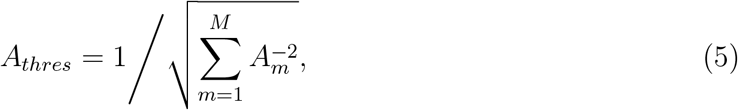

Hearing loss for each frequency depends on *γ*_*ohc*_ and *γ*_*ihc*_, and is defined in *dB* as:

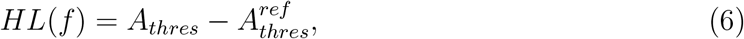

where 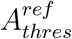 is the reference threshold obtained by setting *γ*_*ohc*_(*x*) = 0.5 and *γ*_*ihc*_(*x*) = 8. These values represent 100% survival of OHCs and IHCs, respectively. The resulting reference threshold in *dB* − *SPL* is shown in Fig. 2, along with the standard threshold given by ISO 226 (ISO, 2003).

**FIG. 2.**
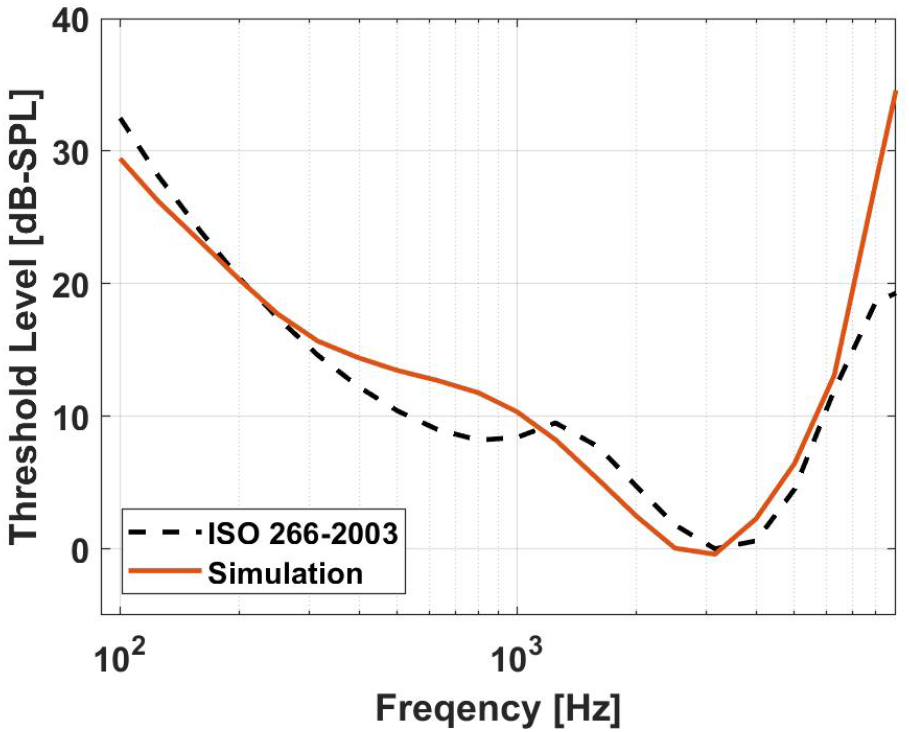
Derivation of auditory threshold (dB SPL) as a function of frequency (red line) compared with the threshold in quiet provided by ISO 226 (dashed black line).

The relation between the hair cells indices (*γ*_*ohc*_ and *γ*_*ihc*_) and the percentage of surviving OHCs and IHCs (*P*_*ohc*_ and *P*_*ihc*_) is given by:

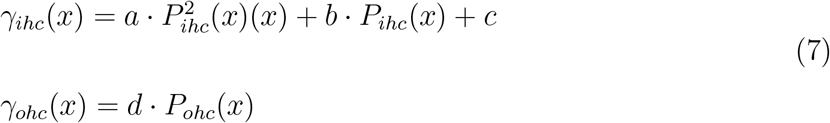

where *a* = 7 *·* 10^−4^; *b* = 0.016; *c* = 3.7; and *d* = 5 *·* 10^−3^.

All computational derivations were implemented using parallel programming written in CUDA for the Nvidia GTX 1080 TI GPU (Furst, 2015; Koral, 2018; Sabo *et al*., 2014).

### B. Algorithm for Estimating Hair Cell Survival from Measured Audiograms

The following algorithm is designed to predict the percentage of surviving hair cells along the cochlear partition based on a given audiogram. The steps of this algorithm are outlined in the flowchart shown in Fig. 3.

**FIG. 3.**
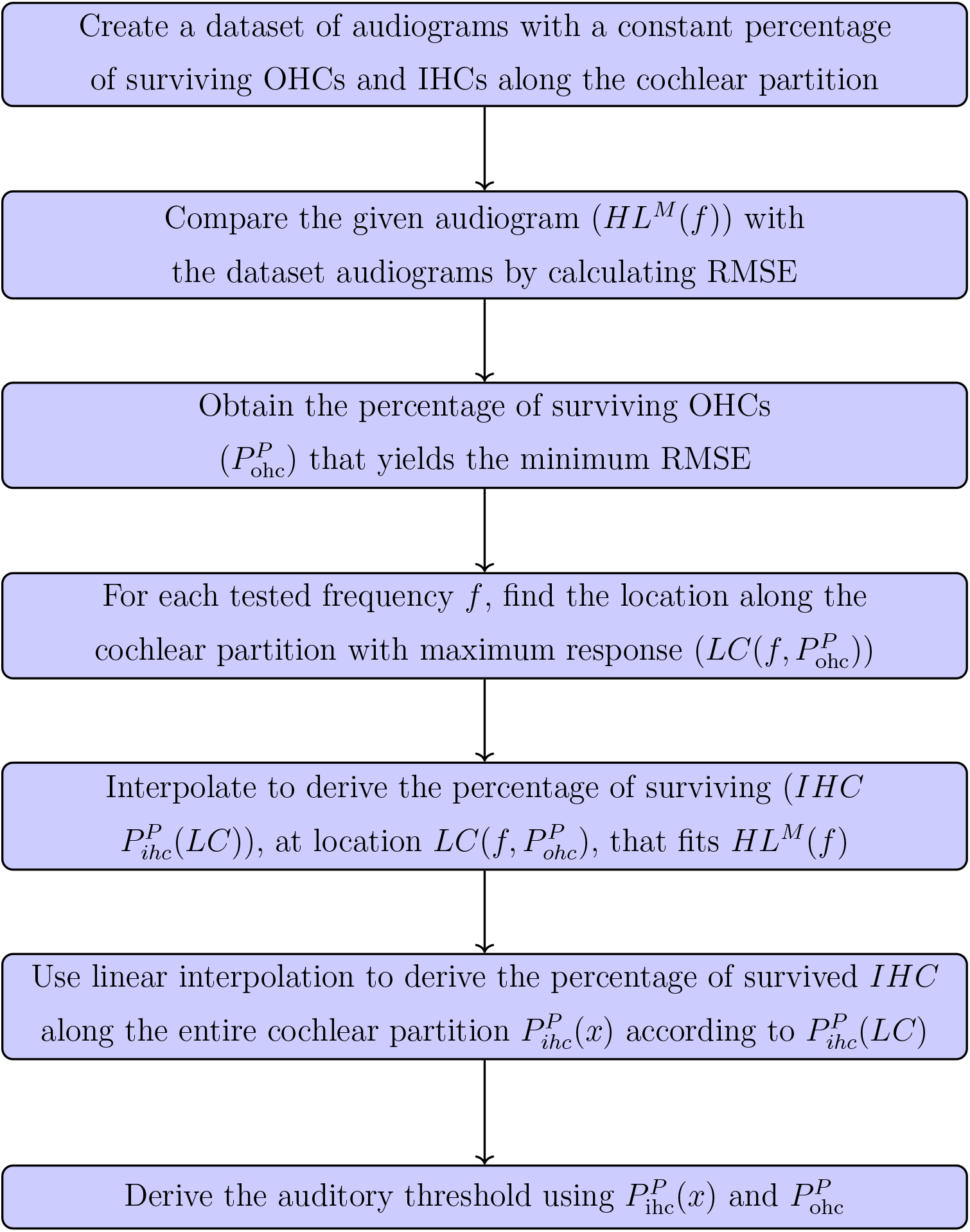
Flowchart of the Algorithm for Estimating Hair Cell Survival given an audiogram *HL*^*M*^(*f*)

The algorithm initiates by creating a dataset of auditory thresholds, derived using constant values of *γ*_*ihc*_ and *γ*_*ohc*_ along the cochlear partition. The resulting dataset is illustrated in Fig. 4, where the hearing level in dB HL (Eq. 6) is plotted as a function of *P*_*ihc*_. Each panel in the figure corresponds to a different input frequency, with colors distinguishing the *P*_*ohc*_ values. Generally, as *P*_*ihc*_ increases, the auditory threshold decreases across all frequencies. However, the degree of this decrease varies; at frequencies above 1000 Hz, it strongly depends on *P*_*ohc*_; at lower frequencies, such as 250 Hz, the impact of *P*_*ohc*_ is minimal.

**FIG. 4.**
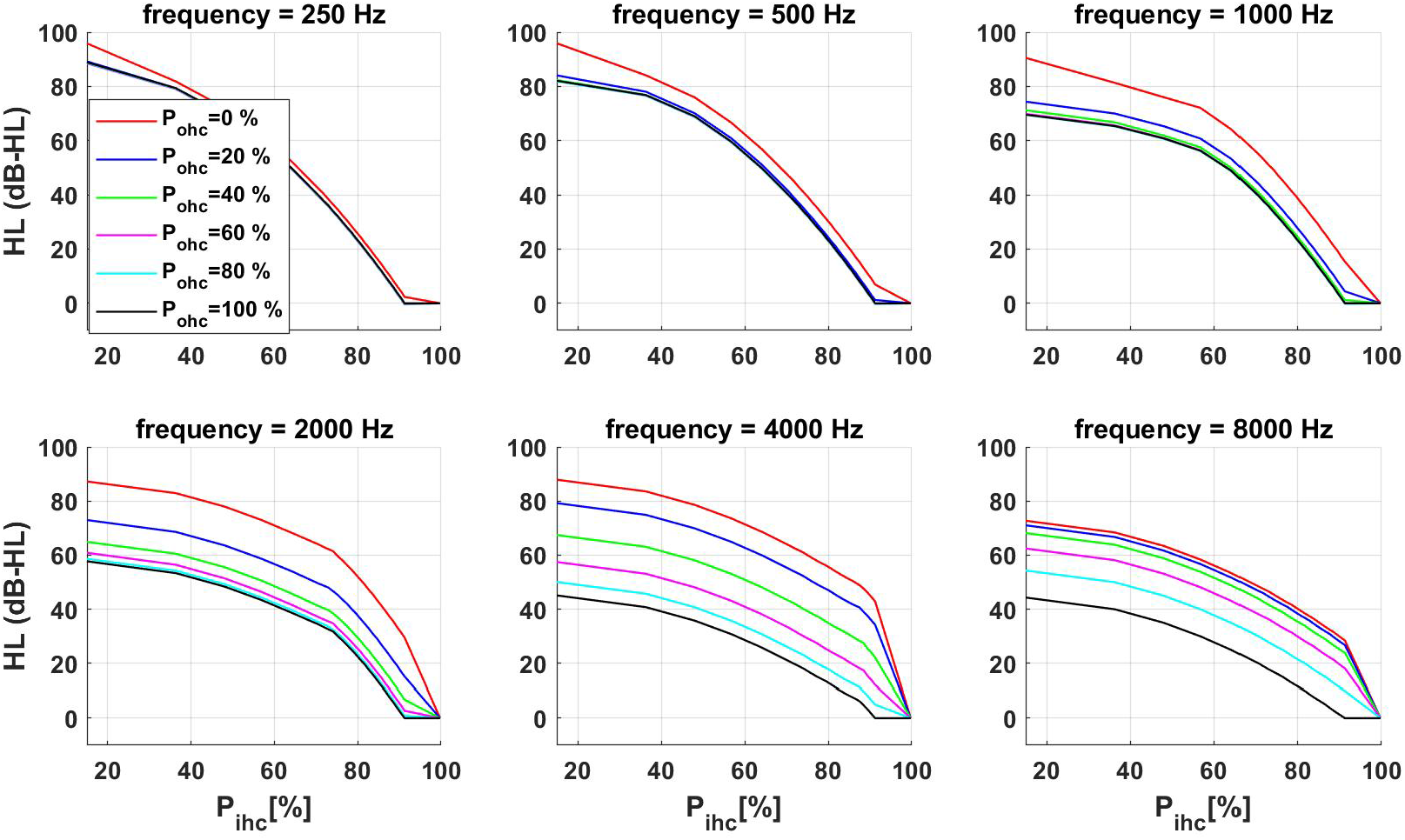
Derived hearing loss as a function of *P*_*ihc*_ for different frequencies and values of *P*_*ohc*_. Each panel represents a different frequency and each color represents a different *P*_*ohc*_.

The algorithm employs the root mean squared error (RMSE) to match measured and calculated hearing levels (HL), defined as:

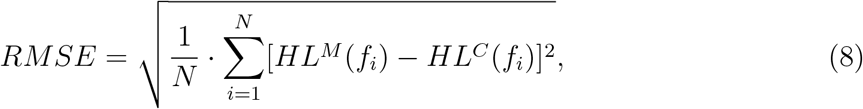

where *HL*^*M*^ and *HL*^*C*^ are the measured and calculated HL, and *N* is the number of the tested frequencies.

For each measured auditory threshold (*HL*^*M*^(*f*)), the *RMSE* is calculated across the entire dataset. The algorithm selects the *P*_*ohc*_ that minimizes the *RMSE* as the predicted survived *OHC* 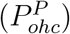:

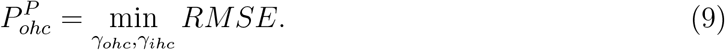

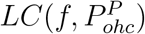 is defined as the cochlear location yielding the maximum BM response for a given frequency *f* and the predicted survived 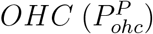. The estimated location is plotted in Fig. 5 as a function of *P*_*ohc*_ for various frequencies. Note that *LC* increases differently for each frequency as a function of *P*_*ohc*_.

**FIG. 5.**
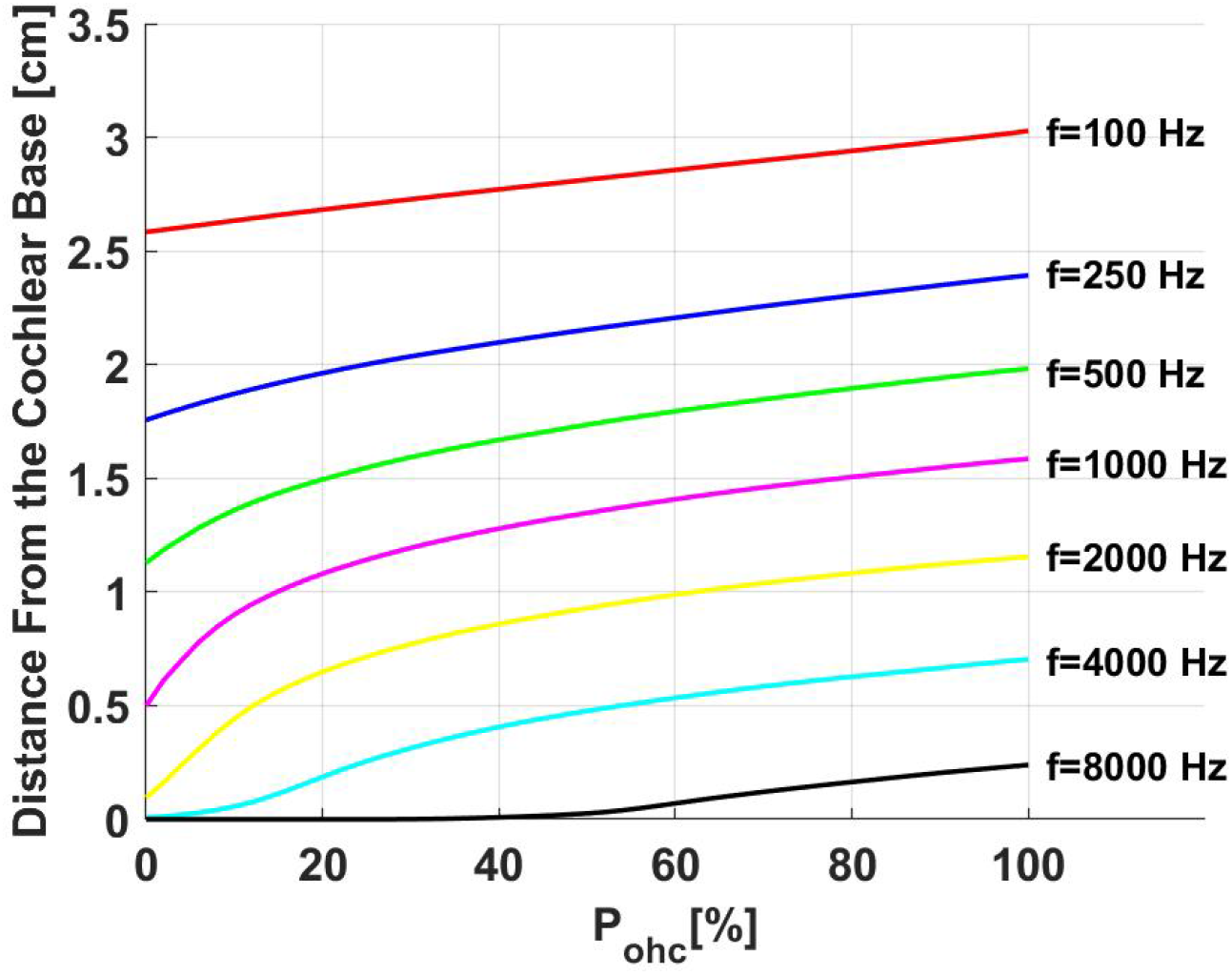
The cochlear position of maximum response as a function of *P*_*ohc*_ for different frequencies

To elucidate the algorithm, we present an illustrative example: an audiogram indicating a hearing loss of 55 dB at a frequency of 2000 Hz. By comparing this audiogram with our dataset, we find OHC survival rate of 40% 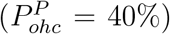. Referring to Fig 5, we can identify the location along the cochlear partition that yields the maximum BM response for a 2000 Hz sine wave. This location is determined to be 0.8 cm from the cochlear base on the y-axis, corresponding to the observed 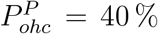. The subsequent step involves calculating the survival percentage of IHC. This calculation is performed using Fig 4, which shows an IHC survival rate of 48% at the 0.8 cm mark 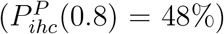. This method is repeated across all frequencies tested in the audiogram, yielding a series of *N* discrete values for 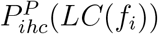, when *i* = 1 : *N*. The final profile of 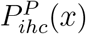 for 0 ≤ *x* ≤ 3.5 *cm* is derived by linear interpolation of these values.

## III. RESULTS

### A. Audiogram Estimation Based on Measured Hair Cells

Wu et al. (Wu *et al*., 2020) presented four typical cases where the audiograms, measured close to death, are displayed alongside postmortem counts of surviving inner hair cells (IHCs) and outer hair cells (OHCs).

Based on the Figure 2 in Wu et al. (Wu *et al*., 2020), the percentages of surviving IHCs and OHCs for four subjects are presented in rows 2 and 3 of Fig.6, indicated by blue circles. The percentage of the surviving hair cells is plotted as a function of the characteristic frequency (*CF*), derived using the Greenwood map (Greenwood, 1990):

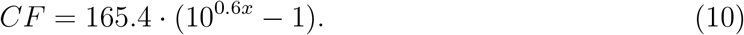

**FIG. 6.**
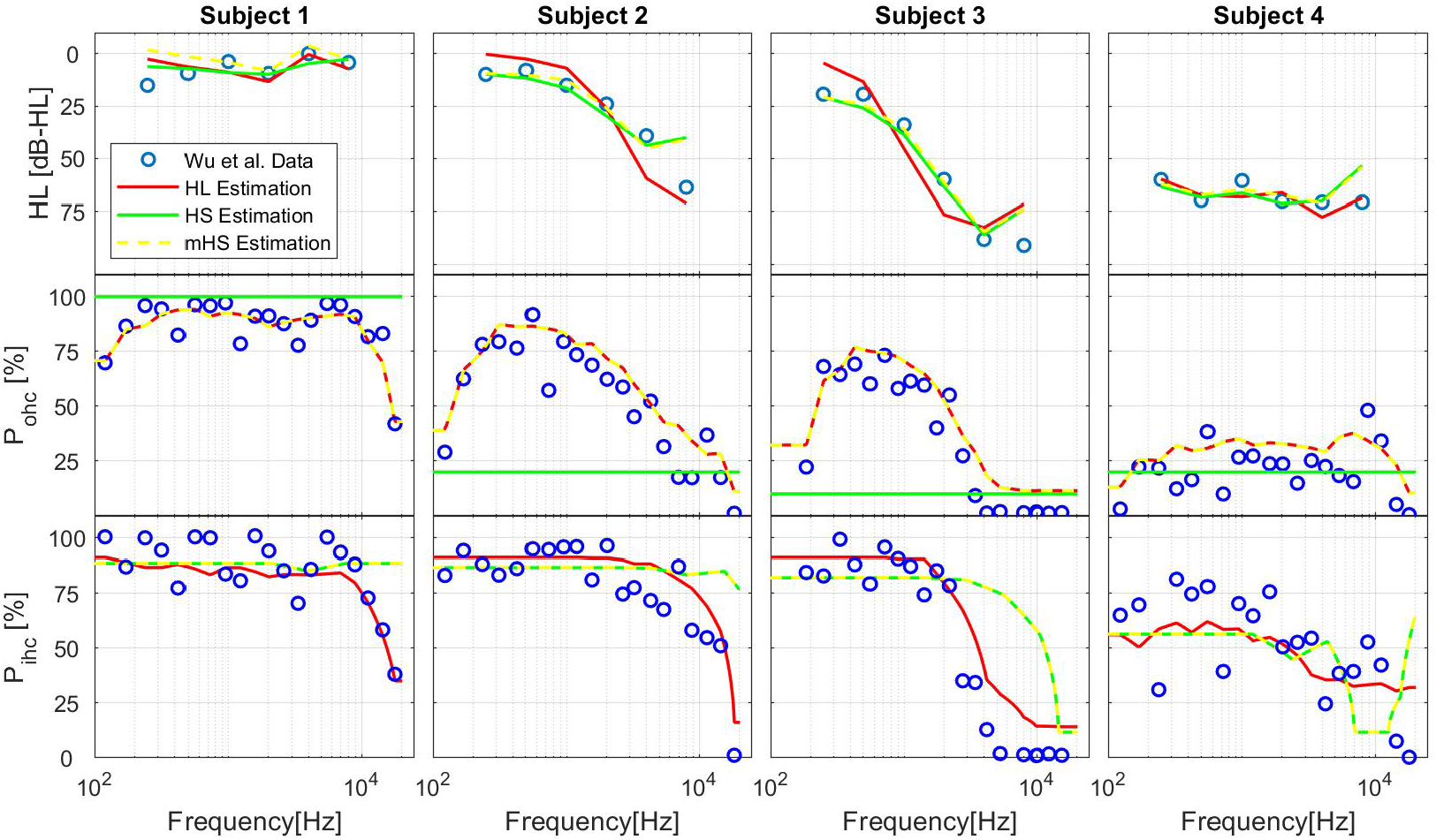
Comparison between data from Wu et al. (blue circles), and the performance of the three algorithms (solid lines): Algorithm 1-red, Algorithm 2-green, and Algorithm 3-yellow. The top row represents the measured and predicted audiograms. The middle and bottom rows represent the measured and predicted percentage of survived OHCs and IHCs, respectively.

To derive the hearing loss (Eq. 6), hair cell indices were obtained by substituting *P*_*ihc*_(*x*) and *P*_*ohc*_(*x*) in Eq. 7. The comparison between the measured data (blue circles) and the estimated auditory thresholds (red lines) is shown in the top row of Fig. 6. To obtain a minimal *RMSE* (Eq. 8), we tested combinations of *P*_*ihc*_+Δ and *P*_*ohc*_+Δ, with −15% ≤ Δ ≤ 15% in the derivation of *HL*^*C*^. The resulting values for *P*_*ihc*_, *P*_*ohc*_, and *HL*^*C*^ are depicted in Fig. 6 by red lines, labeled as hearing level estimation (“*HL* Estimation”). The figure clearly shows that for all four subjects, the model accurately predicted the auditory thresholds based on the measured surviving hair cells. A statistical t-test showed no significant difference between the measured and calculated audiograms (*p* > 0.7) for all subjects.

### B. Prediction of the Survived Hair Cells Based on the Auditory Threshold

This section explores whether the auditory threshold alone can predict the percentage of surviving hair cells along the cochlear partition. The calculated *HL*^*C*^ and the predicted percentage of the hair cell survival for the four subjects from Wu et al. data (Wu *et al*., 2020) are represented by the green solid lines in Fig. 6, labeled as Hair Cell Estimation (“*HS* Estimation”).

The comparison between the measured surviving IHCs and the estimated ones is displayed in the lower row of Fig. 6. No significant difference was found between them for all subjects (*p* > 0.05).

The algorithm described in SEC. II B yields a constant *P*_*ohc*_ along the cochlear partition, as shown by the green horizontal lines in the middle row of Fig. 6. For subject 4, whose surviving OHCs are almost constant along most of the cochlear partition, the predicted surviving OHCs percentage closely matches the measured one (*p* = 0.8). For subjects 2 and 3, whose *P*_*ohc*_ changes significantly along the cochlear partition, the prediction is adequate for *f* > 2500 *Hz* (*P* > 0.1). For subject 1, the algorithm predicts *P*_*ohc*_ = 100%, while measurements show lower values, resulting in a difference between the measured and predicted audiograms (*p <* 0.05).

The calculated audiogram with the predicted *P*_*ihc*_ and constant *P*_*ohc*_ are shown in the top row of Fig. 6. No significant difference was found between the measured and calculated audiograms (*p* > 0.6).

Since different profiles of OHCs along the cochlear partition can yield similar audiograms (Cohen and Furst, 2004; Faran and Furst, 2023), we tested the algorithm’s predictions using the measured *P*_*ohc*_ and estimated *P*_*ihc*_. The results are represented by the yellow lines in Fig. 6, labeled as modified hair cell estimation (“*mHS* Estimation”). No significant differences were found between the audiograms derived using the constant *P*_*ohc*_ (green lines) and the variable *P*_*ohc*_ (yellow lines), with a t-test yielded *p* = 0.04 for subject 1, and *p* > 0.8 for subjects 2, 3, and 4. Comparison between the measured audiograms and predictions from Algorithm 3 revealed no significant difference for all subjects (*p* > 0.1).

## IV. DISCUSSION

The relation between audiograms and hair cell loss was demonstrated by applying a cochlear computational model to data from Wu et al. (Wu *et al*., 2020). We have shown that the measured audiogram can be predicted using the surviving IHCs and OHCs. Additionally, we introduced an algorithm that predicts the surviving hair cells along the cochlear partition based on the measured audiogram. The algorithm predicts a constant percentage of OHCs along the cochlear partition, whereas the measurements reveal a variable distribution of surviving OHCs. The predictions accurately match only the surviving OHCs at the basal part of the cochlea. However, we have demonstrated that a similar audiogram can be obtained using either the constant predicted OHC survival percentage or the variable mea-sured percentage of OHCs. This theoretical result aligns with experimental observations in mice, which suggest that OHCs contribute less to cochlear amplification at low frequencies (Liberman *et al*., 2002). It also corresponds with human studies indicating a U-shaped audiogram with a maximum threshold shift at mid-high frequencies (2-4 kHz) due to noise exposure in young adults (Hannula *et al*., 2011; Moore, 2007; Saunders *et al*., 1985). In the prediction algorithm (*HS* Estimation), the percentage of IHCs is determined based on the location of the maximum response for each frequency. As illustrated in Fig. 5, this location depends on the percentage of surviving OHCs. This theoretical prediction is consistent with animal data showing that damage to OHCs causes a shift in the characteristic frequency (CF) location for both auditory nerve responses (Liberman, 1984) and basilar membrane motion (Ruggero and Rich, 1991).

The prediction algorithm appears adequate for IHC loss but cannot precisely predict OHC loss along the cochlear partition. As demonstrated, both constant and variable OHC loss can produce audiograms similar to the measured data. It is most likely that other auditory properties, such as psychophysical tuning curves or auditory masking, are more sensitive to the distribution of surviving OHCs.

The present study tested only measurements from four subjects; more data should be collected, including other types of damage, such as malfunctioning of the stria vascularis, which may also affect the audiogram (Schmiedt, 1996; Vaden *et al*., 2022). Nonetheless, this study demonstrates the potential of using cochlear modeling to predict cochlear damage from audiograms.

## APPENDIX A COMPUTATIONAL MODEL

This section covers the cochlear model, where the parameters and constants of the model are summarized in Appendix. B.

### a. Deriving the Basilar Membrane Motion

By applying the conservation of mass and the dynamics of deformable bodies principles (Cohen and Furst, 2004), the partial differential equation for the pressure difference across the cochlear partition, *P*, was obtained by:

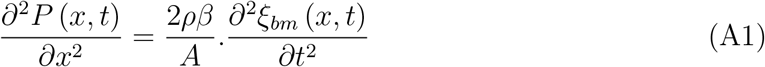

where *x* is the cochlear position, *t* is the time, *ξ*_*bm*_ is the basilar membrane (BM) displacement, *A* is the cross-sectional area of both the Scala Tympani and the Scala Vestibuli, *β* is the BM width, and *ρ* is the density of the fluid in both Scala Vestibule and Scala Tympani.

The OHCs lay on the BM, and their upper part is embedded in the tectorial membrane (TM). The relation between the BM’s, TM’s, and OHCs’ pressures (denoted as *P*_*bm*_, *P*_*tm*_ and *P*_*ohc*_ respectively) can be described as:

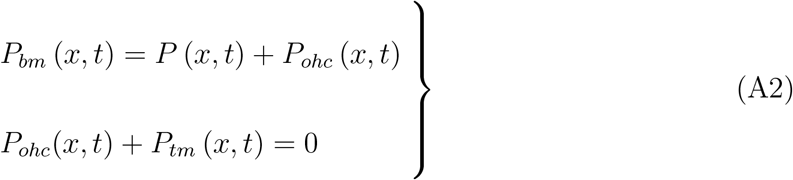

The mechanical properties of both BM and TM were simulated as second-order oscillators:

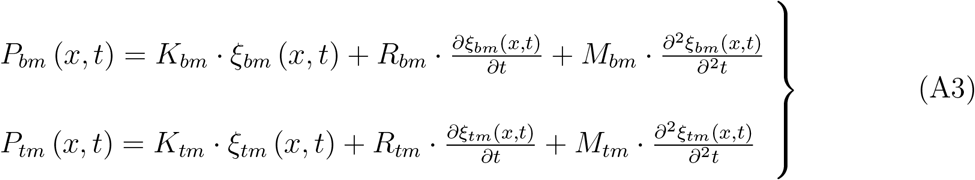

where *ξ*_*bm*_, *ξ*_*tm*_, *K*_*bm*_, *K*_*tm*_, *M*_*bm*_, *M*_*tm*_, *R*_*bm*_, *R*_*tm*_ are the displacement, effective stiffness, mass per unit area, and resistance of the BM and TM respectively.

Since the OHCs are placed between the BM and TM membranes, their displacements *ξ*_*ohc*_ yielded the following equation:

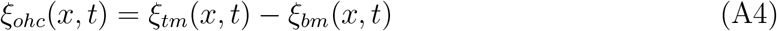

An increase in the OHC’s receptor potential, *ψ*, causes a decrease in their length, which in turn enhances the movement of the BM (Brownell *et al*., 1985). The length of the OHC, *δ*_*ohc*_, can be described as a sigmoid function of *ψ* (He and Dallos, 2000; Kakehata and Santos-Sacchi, 1995),

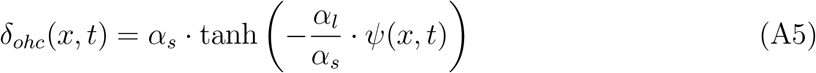

where *α*_*s*_ and *α*_*l*_ are constants.

The OHC’s electrical potential *ψ* was obtained by solving the equivalent electrical circuit (Barzelay and Furst, 2011; Cohen and Furst, 2004), which provided the following equation:

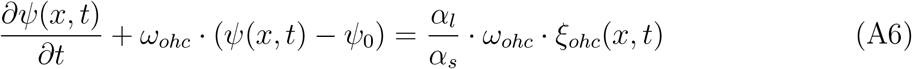

where *ω*_*ohc*_ and *ψ*_0_ denote the OHC’s cutoff frequency and the perilymph resting potential, respectively.

The OHC’s pressure obtained from the elastic properties of the OHCs (Cohen and Furst, 2004). As a result, *P*_*ohc*_ can be calculated using the following equations:

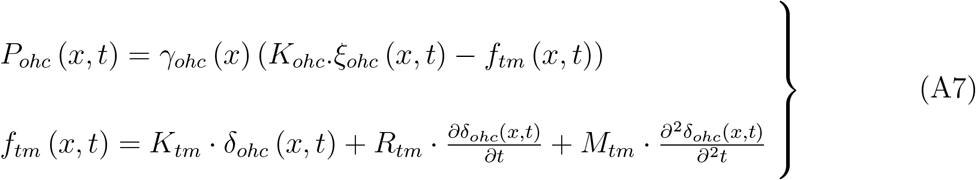

where *K*_*ohc*_ is the OHC’s stiffness, and *γ*_*ohc*_(*x*) represents the effective distribution of OHCs along the cochlear partition. A cochlea with no active OHC is obtained by *γ*_*ohc*_(*x*) = 0. *γ*_*ohc*_(*x*) = 0.5 yields a healthy cochlea that most accurately matches physiological results (Furst, 2015).

The ear model was solved by applying initial and boundary conditions. The initial conditions were all set to zero. The boundary conditions are:

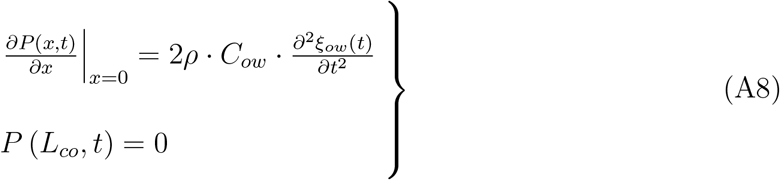

where *L*_*co*_ is the cochlear length, *ξ*_*ow*_ is the Oval Window displacement, and *C*_*ow*_ is the coupling factor of the Oval Window to the perilymph.

In order to obtain the *ξ*_*ow*_, the middle and outer ear model expressed by the following differential equation (Talmadge *et al*., 1998), was applied:

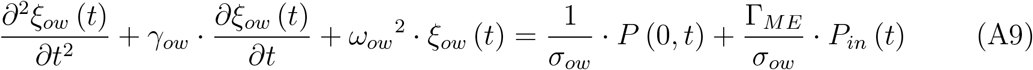

where *ω*_*ow*_ is the oval window resonance frequency, Γ_*ME*_ is the mechanical gain of the ossicles, *γ*_*ow*_ is the oval window resistance, *σ*_*ow*_ is the oval window areal density, and *P*_*in*_(*t*) is the input acoustic stimulus in *Pa*.

The initial conditions were all set to zero, specifically :

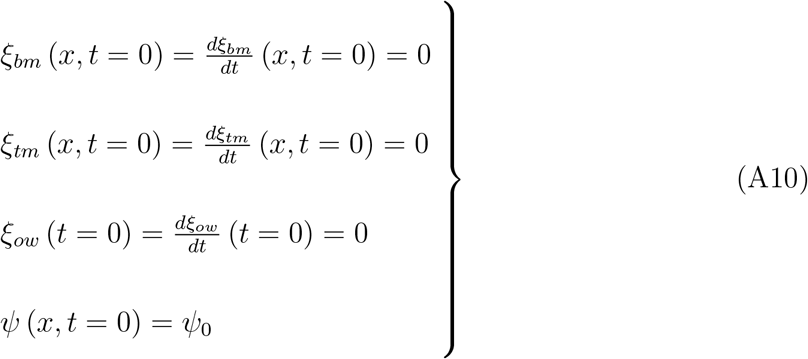

#### b. Auditory Nerves’ Instantaneous Rates

The BM motion is transformed into neural spikes of the ANs by the IHCs. The BM displacement stimulates the IHC cilia to move, therefore its displacement *ξ*_*ihc*_ can be approximated as:

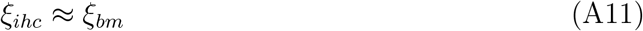

The IHCs’ transduction abilities were modeled as a nonlinear system that combines AC and DC components (Sumner *et al*., 2002; Zilany *et al*., 2009). The IHC’s membrane electrical potential, *ψ*_*ihc*_, was obtained by:

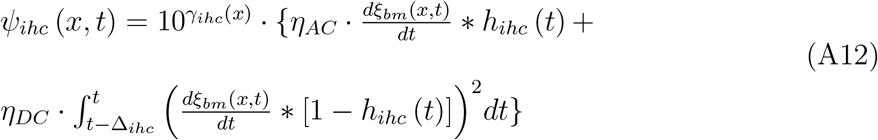

where *h*_*ihc*_(*t*) is a low-pass filter that represents the IHCs impulse response, *η*_*AC*_, *η*_*DC*_, and Δ_*ihc*_ are constants. *γ*_*ihc*_(*x*) represents the IHC efficiency index. For normal cochlea, *γ*_*ihc*_(*x*) = 8 was found to match experimental data (Furst, 2015).

One possible approach to describe the stochastic properties of neural activity relates to the spikes train as a non-homogeneous Poisson point process, whose IR depends on the input stimuli (Gray, 1967; Rieke *et al*., 1999; Rodieck *et al*., 1962). If *λ*_*AN*_ represents the sensation, and *ψ*_*ihc*_ represents the stimulus, then *λ*_*AN*_ can be obtained by (Furst, 2015),

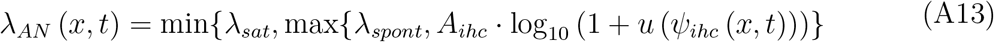

where *λ*_*spont*_ and *λ*_*sat*_ are the spontaneous and saturation ANs’ IRs respectively, *u*(*t*) is a step function and *A*_*ihc*_ is the AN’s coupling constant.

## APPENDIX B LIST OF MODEL PARAMETERS

**TABLE I.**
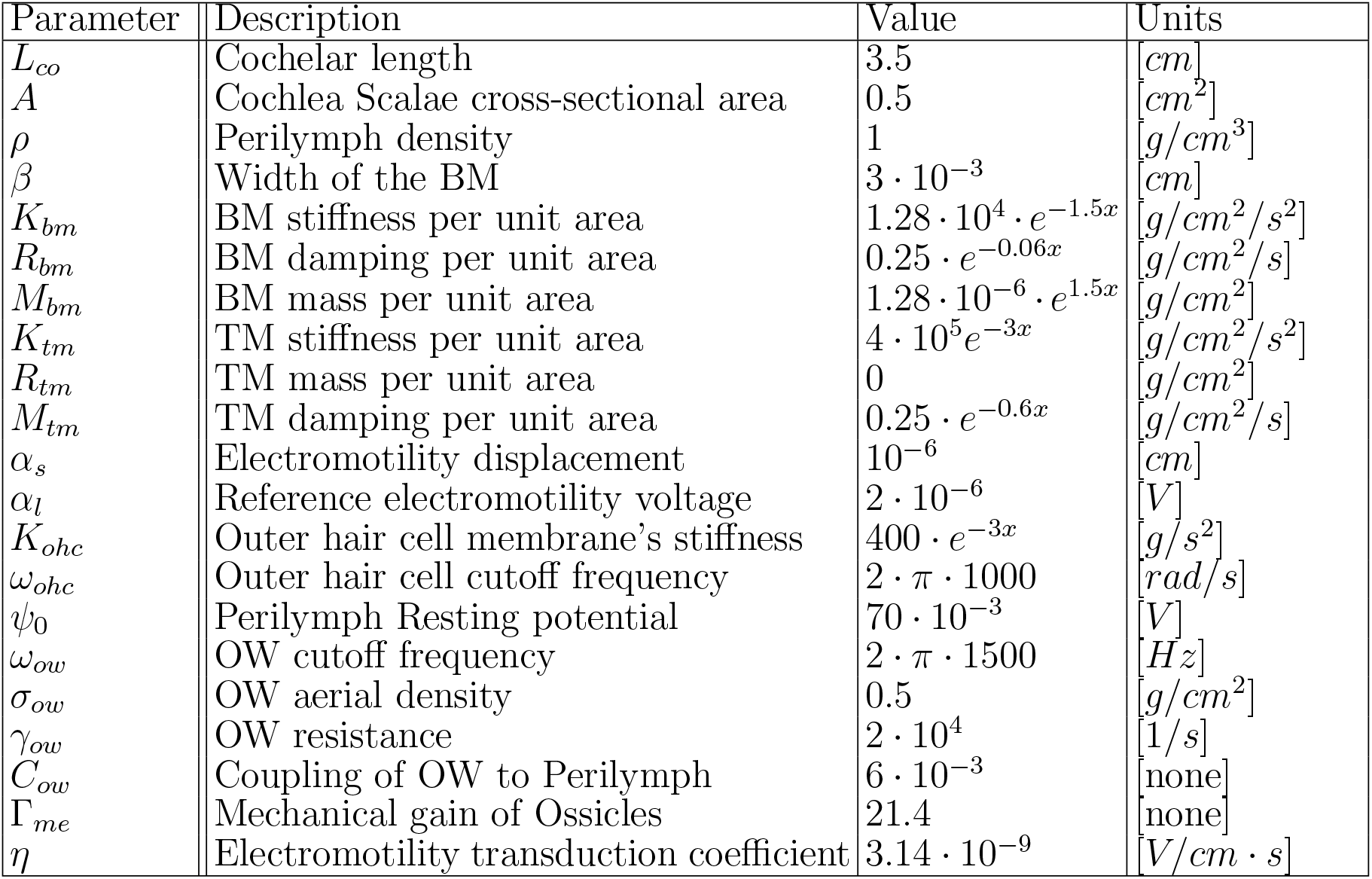
BM parameters.

**TABLE II.** IHC parameters.

| Parameter | Description | Value | Units |
| --- | --- | --- | --- |
| $\eta_{AC}$ | IHC AC coupling | 1 | [V/s/cm] |
| $\eta_{DC}$ | IHC DC coupling | 100 | [V/cm] |
| $\Delta_{ihc}$ | IHC integration time | $2 \cdot 10^{-3}$ | [sec] |
| $A_{ihc}$ | ANF coupling | 1 | [spikes/s/V] |
| $\lambda_{sat}$ | Saturation rate | 500 | [spikes/s] |
| $\lambda_{spon}$ | Spontaneous rate | 0.1 | [spikes/s] |
| $f_{co}(h_{ihc})$ | The cut-off frequency of the IHC | 800 | [Hz] |

